# RNA regulates cohesiveness and porosity of a biological condensate

**DOI:** 10.1101/2024.01.09.574811

**Authors:** Han-Yi Chou, Aleksei Aksimentiev

## Abstract

Biological condensates have emerged as key elements of a biological cell function, concentrating disparate biomolecules to accomplish specific biological tasks. RNA was identified as a key ingredient of such condensates, however, its effect on the physical properties of the condensate was found to depend on the condensate’s composition while its effect on the microstructure has remained elusive. Here, we characterize the physical properties and the microstructure of a protein–RNA condensate by means of large-scale coarse-grained (CG) molecular dynamics simulations. By developing a custom CG model of RNA compatible with a popular CG model of proteins, we systematically investigate the structural, thermodynamic, and kinetic properties of condensate droplets containing thousands of individual protein and RNA molecules over a range of temperatures. While we find RNA to increase the condensate’s cohesiveness, its effect on the condensate’s fluidity is more nuanced with longer molecules compacting the condensate and making it less fluid. We show that a biological condensate has a sponge-like morphology of interconnected channels of size that increases with temperature and decreases in the presence of RNA. Our results suggest that longer RNA form a dynamic scaffold within a condensate, regulating not only its fluidity but also permeability to intruder molecules.

---

Ribonucleoprotein granules—membraneless assemblies formed by various RNA-binding proteins and nucleic acids—have recently become a subject of intense scientific inquiry because of their newly discovered roles in diverse aspects of cell biology, including gene expression, regulation, cell signaling and RNA concentration buffering ^1–7^. Although such membraneless organelles can be formed by a variety of seemingly disparate biomolecular species, their behavior is believed to be described by a common physical mechanism, the so-called liquidliquid phase separation (LLPS), where a condensed, liquid-like droplet containing the proteins and the nucleic acids coexist with a much more dilute phase of the same molecules in a dynamic equilibrium^8–10^.

Previous experimental work revealed that many RNA-binding proteins, such as DEAD-box helicase LAF-1 and fused in sarcoma (FUS), display LLPS *in vivo* and *in vitro* ^8,11–14^. The studies have also found RNA to modulate the structural and dynamic properties of liquid droplets formed by such RNA-binding proteins in a non-trivial manner. Thus, the presence of long RNA strands was found to enhance the condensation propensity of FUS ^15^, whereas excessive amounts of RNA can lead to a dissolution of FUS condensates^16^ in a phenomenon known as reentrant phase behavior. The droplets formed by LAF-1 become more fluidic when RNA is added^13^. The droplets formed by FUS, on the other hand, can become less viscous in the presence of short RNA molecules and more viscous in the presence of long RNA molecules ^17^. The microscopic mechanisms by which RNA modulates the physical properties of the condensates largely remain elusive.

Despite tremendous advances in the experimental methodologies that now permit accurate characterization of the structural and dynamic properties of single molecules in solution, similar characterization of biomolecules residing within the condensed droplets remains challenging^18^. Obtaining the information inaccessible to experimental probes is, however, possible using theoretical and computations approaches^19^. Self consistent field theory^20^, lattice protein models^21–23^ and various molecular dynamics (MD) approaches^24^ have been widely used to decipher the mechanisms underlying LLPS. MD studies of biomolecular interactions enabling the LLPS phenomena differ in their resolution from explicit solvent all-atom models^25–28^ to coarse-grained models that in turn differ from one another by the parameterization of the intermolecular interactions and their treatment of solvent^29–31^. Among the most popular one-bead-per-amino-acid models used for MD simulations of biomolecular condensates are the Kim-Hummer (KH) model^32–34^, the HPS model^33^ and its variants^35–37^, the AWSEM-IDP model^38^, and more specialized models targeting a particular type of disordered proteins^39^. We note, however, that the majority of the CG MD simulations to date have focused on characterization of the protein components of the condensates, in part because of the lack of CG models of RNA compatible with the existing CG models of intrinsically disordered proteins. Addressing that need, a coarse-grained model of RNA was recently developed^30,40^ to match the parameterization framework of the HPS model and applied to investigate the properties of protein–RNA condensats^41^. A systematic approach to obtaining the interaction potential between the nucleic acid components of a biomolecular condensates and their protein counterparts within the framework of an arbitrary CG model remain a subject of ongoing research^42^.

Here, we describe a heuristic yet efficient approach to constructing and calibrating a CG model of RNA that can be generalized to an arbitrary implicit-solvent CG framework. Unlike the previous sophisticated models that incorporate the effect of nucleotide orientation and stacking^43–45^, we employ a much simpler, one-bead-per-nucleotide model of RNA that nevertheless accurately reproduces its polymer physics properties, quantitatively captures the strength of specific interactions with the condensate’s proteins and reproduces the nontrivial behavior of RNA–peptide mixtures, i.e., their reentrant phase behavior. We then use the resulting RNA model to investigate how the length and the concentration of RNA modulates the structure and dynamics of a FUS condensate droplet. We show that the presence of RNA makes the FUS condensate more cohesive, more viscous and less accessible to small molecules by shrinking the size of the passages that perforate the interior of the condensate.

## Results

### CG model of RNA for MD simulations of FUS condensates

We used a top-down approach to derive a one-bead-per-nuleotide CG model of unstructured RNA that is compatible with the KH model^32^ of disordered proteins, Fig. 1a. First, we used small angle X-ray scattering (SAXS) data to determine the bonded and non-bonded parameters describing the conformations of an RNA molecule in isolation. Following that, we determined non-bonded parameters describing the interactions between RNA and FUS, assuming that only the structured domains of FUS bind RNA.

**Figure 1:**
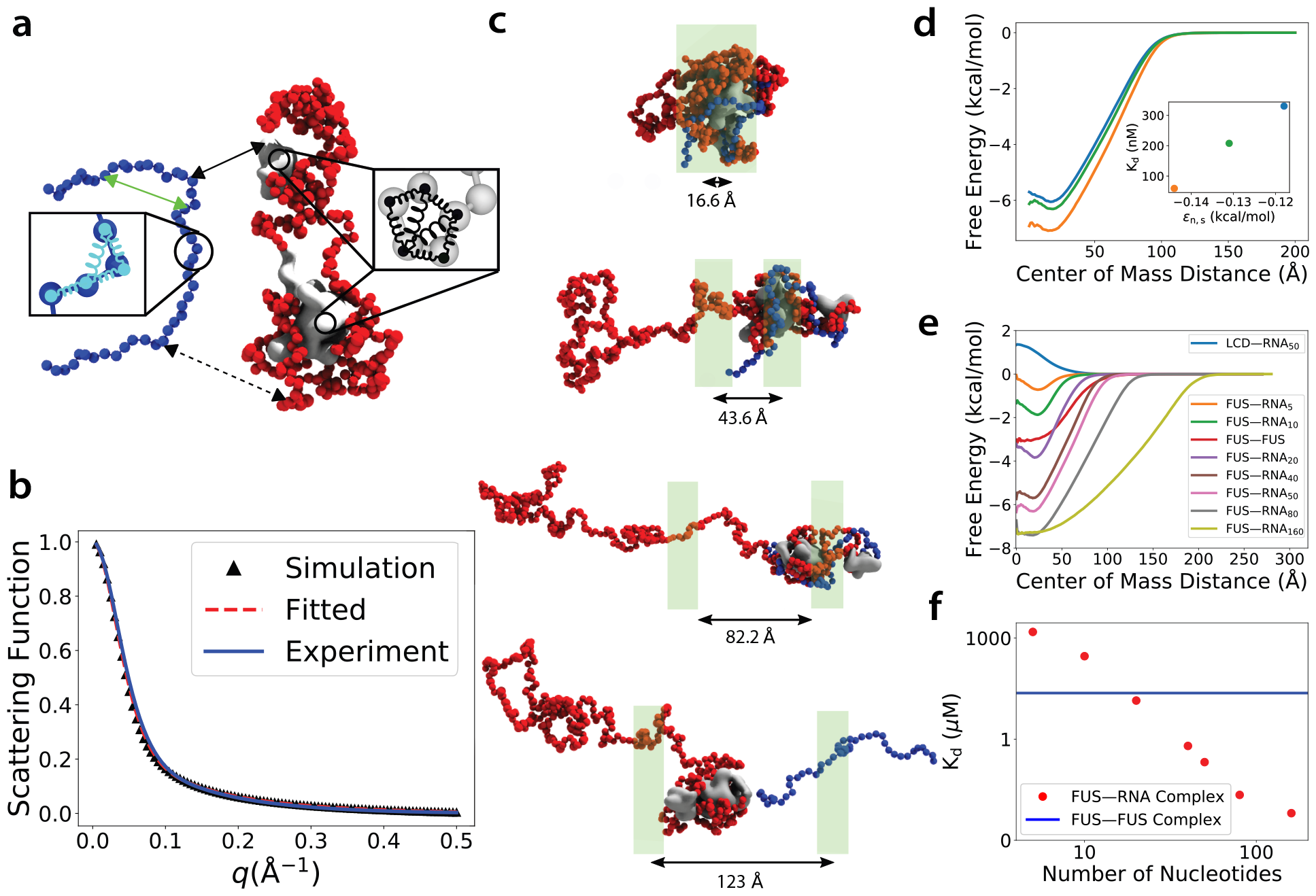
Coarse-grained model of RNA for MD simulations of FUS–RNA condensates. **a**, Schematics of the model’s parameterization. An unstructured RNA stand (blue) is represented as a collection of charged beads connected via bond and angle harmonic potentials. A full length FUS protein (red) is represented at one-bead-per-amino resolution using a previously described model^32^. The conformation of the two structured (silver) domains are enforced using a network of harmonic restraints. The model is parametrized to reproduce the interactions within the unstructured RNA (represented by green arrow), and between the RNA and the structured and the unstructured parts of the FUS protein (solid and dashed black arrows, respectively). **b**, Parameterization of the RNA model against SAXS data^46^. **c**, Adaptive biasing force (ABF) calculations of the free-energy of the FUS–RNA interaction. The four snapshots depict representative conformations of the simulation system at four sampling windows. The instantaneous distance between the centers of mass of the FUS and the RNA, which is the ABF reaction coordinate, is specified for each snapshot below the black arrow. The green rectangles represent the range (to scale) of the reaction coordinate values explored within each window. **d**, The potential of mean force (PMF) between a full-length FUS and a 50-nucleotide ssRNA molecule as a functions of the *E*_n,s_ parameter that prescribes the strength of the interactions between the structured residues of FUS and the RNA nucleotides. The PMFs for the three values of *E*_n,s_ and the corresponding *K*_*d*_ values (inset) are shown using matching colors. **e**, FUS–FUS and FUS–RNA PMFs for RNA of different lengths. The interaction between the low complexity domain of FUS and the RNA is repulsive by the very definition of the model. **f**, The dissociation constants of a FUS–RNA complex as a function of the RNA length. The blue line indicates the dissociation constants of a binary FUS–FUS complex.

According to SAXS measurements^46^, the contour length, *l*_c_, of a poly(U)_40_ ssRNA is 196.4 Å, whereas its persistent length, *l*_p_ = 21.0 Å. In our CG model of ssRNA, two beads representing a pair of neighboring nucleotides are prescribed an equilibrium bond length *l*_0_ whereas any three consecutive beads are prescribed an equilibrium valence angle *θ*_0_. Following the approach described in Ref. 47, we determined the average bond length *l*_0_ = 5.036 Å using the relation *l*_*c*_ = (*N* − 1)*l*_0_, where *N* is the number of nucleotides within the RNA. Similarly, we determined the equilibrium valence angle value *θ*_0_ = 141.8 degrees using the relation^47^ 2*l*_*p*_ = *l*_0_(1 − cos *θ*_0_)*/*(1 + cos *θ*_0_).

We validated our parameterization of the RNA model by simulated a 40-nucleotide RNA strand at 295.12 K for 4 *µ*s, discarding the first 500 ns, and using the remaining trajectory to compute the structure factor

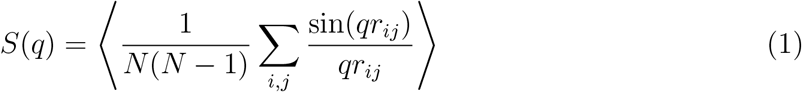

as a function of the scatting vector, *q*, which was then used to determine the contour and persistent lengths of the polymer. In doing so, we were aiming to reproduce directly the properties measured by the SASX method. The simulated *S*(*q*) function was found to be in good agreement with the one determined by experiment, Fig. 1b. Finally, the simulated *S*(*q*) was fitted to a semi-analytical function (based on a worm-like-chain model) in high-*q* region (from 0.1 to 0.5) to independently obtain the contour length and the persistent length in accordance with the procedure used to analyze the experimental data^46^. The contour and persistent lengths determined by fitting our simulated scattering function were 197.8 Å and 23.4 Å respectively, in good agreement with the experimentally derived values ^46^.

Informed by experiments showing high affinity of the RBM and Zinc finger domains of FUS to RNA^48^, and by a study demonstrating that electrostatic interactions are sufficient to ensure high affinity binding of intrinsically disorder proteins^49^, we parametrized the FUS–RNA interactions assuming attractive forces between residues forming the structured domains of FUS whereas the interactions between RNA and disordered residues were assumed to be weak and repulsive. Thus, we assigned a small, positive value of 0.001 kcal/mol to the energy scale parameter *ϵ*_n,d_ that describes the interactions between an RNA nucleotide and an amino acid from disordered regions of FUS. We then determined the energy scale parameter *ϵ*_n,s_ describing the interaction of an RNA nucleotide and an amino acid from the structural domains of FUS by matching the experimentally determined dissociation constant, *K*_*d*_.

Following the derivation presented elsewhere^49,50^, the *K*_*d*_ of a FUS–RNA complex can be expressed in terms of the effective interaction potential, *F*_eff_, between the FUS and the RNA,

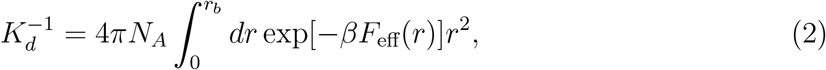

where *r* is the FUS–RNA center-of-the-mass distance, *N*_*A*_ is the Avogadro number and *r*_*b*_ is the radius of the binding complex which is defined as the maximum distance when *F*_eff_ become positive^49^. We determined the effective interaction potential, *F*_eff_, by using the adaptive bias force (ABF) method^51^ to compute the potential of mean force (PMF) between the full length FUS protein and an RNA molecule; Fig. 1c schematically illustrates the ABF procedure.

We performed our calibration simulations using a 50-nucleotide RNA molecule, which has an experimentally determined *K*_*d*_ value for interaction with FUS of 178 nM^52^. Since the dissociation constant is a monotonic function of *ϵ*_*ns*_, we searched the optimal value for ϵ_n,s_ using a binary-search method. Figure 1d shows the resulting PMFs for three values of the ϵ_n,s_ parameter, whereas the inset plots the *K*_*d*_ value as a function of ϵ_n,s_. Based on the results of these simulations, ϵ_n,s_ was set to −0.131 kcal/mol.

Having the FUS–RNA interactions parametrized, we determined the dependence of the effective interaction potential between FUS and RNA, Fig. 1e, and of the dissociation constant, Fig. 1f, on the length of the RNA molecule. Increasing the RNA length is seen to decrease the dissociation constant exponentially, until approximately 90 nucleotides, after which the dependence becomes less pronounced. For reference, we also determined the effective potential between a 50-nucleotide RNA and the low complexity domain (LCD, residue 1 to 163) of FUS, Fig. 1e, finding the expected short-range repulsion. In contrast, a binary FUS–FUS interaction was attractive, with a *K*_*d*_ value comparable to that between a single FUS and a 20-nucleotide RNA, Fig. 1f.

### The CG model recapitulates reentrant phase behavior of a model condensate

As an independent test of our model, we investigated the model’s ability to reproduce reentrant phase behavior experimentally observed^16,53^ in mixtures of RNA and short cationic peptides, abbreviated as RP_3_ peptides^54^. At a fixed concentration of RP_3_, the reentrant phase behavior is manifested by first appearance and then dissolution of the condensate phase as the concentration of RNA molecules is increased, Fig. 2a. Reproducing the experimental conditions, we simulated a system containing 64 RP_3_ peptides having the {RRASL}_3_ amino acid sequence and either 2, 4, 8, 16 or 24 copies of RNA_40_ molecules, which corresponds to the RNA-to-RP_3_ weight ratio of 0.23, 0.48, 0.96, 1.92 or 2.88, respectively. The simulation unit sell was 810.4 × 810.4 × 810.4 Å^3^ and the peptide concentration was 200 *µ*M, matching the conditions of the corresponding experiment^16^. Each simulation was run for Figure 2b shown the initial (top row) and the final (bottom row) configurations of the mixtures containing 2, 8 and 24 RNA_40_ molecules, whereas Movie 1, 2 and 3 illustrate the respective MD trajectories. Qualitatively, our simulations appear to reproduce the reentrant phase behavior: no phase separation is observed at the lowest and the highest concentration of RNA. To characterize the phase separation process quantitatively, we defined a partition coefficient for the largest cluster as the ratio of the number of molecules forming the largest cluster to the total number of molecules in the simulation system. Plotting that partition coefficient as a function of time, Fig. 2c, shows that each system reached equilibrium within the time scale of our simulations. The plot of the partition coefficient averaged over the last 3 *µ*s of each simulation as a function of the RNA-to-RP_3_ weight ration, Fig. 2d, clearly shows the reentrant phase behavior, where droplet formation is suppressed when few RNA molecular are present, the tendency for phase separation increases with the weight ratio, dropping abruptly as the weight ratio exceeds a factor of 3, indicating a disappearance of the condensed phase. Comparing to the behavior observed in experiment, Fig. 2d, we conclude that our CG model reproduces the reentrant phase behavior of an RP_3_–RNA_40_ mixture.

**Figure 2:**
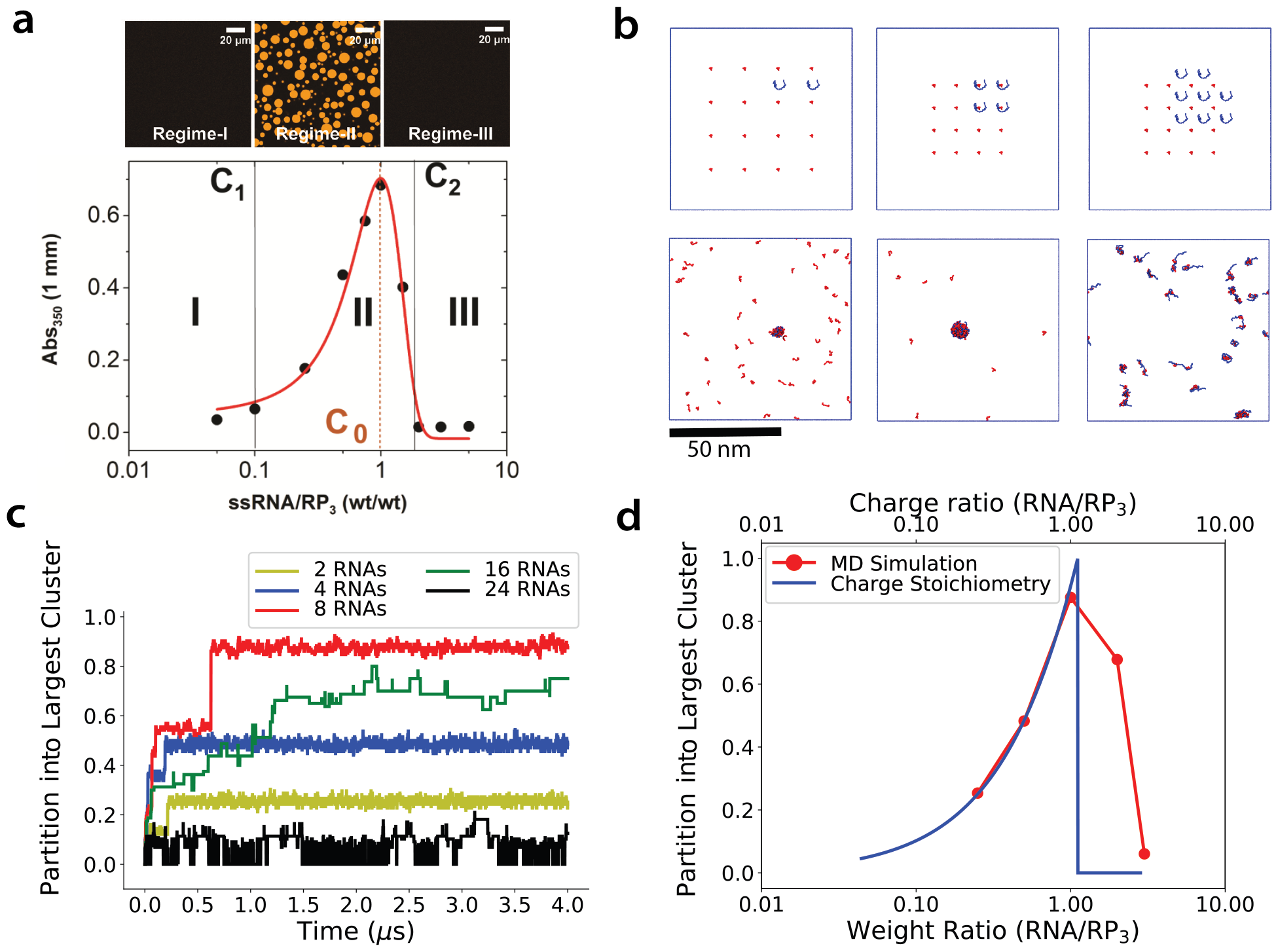
RNA model reproduces reentrant phase behavior. **a**, Reentrant phase behavior of a RP_3_ – RNA_40_ mixture determined experimentally by Banerjee et. al. ^16^. The mixture’s turbidity is plotted as a function of the RNA-to-protein weight ratio, while the peptide’s concentration remains constant at 200 *µ*M. The images illustrates representative morphology of the mixture corresponding to the regions defined in the turbidity plot. Adapted with permission from Ref. 16. Copyright 2017, John Wiley and Sons Angewandte Chemie International Edition. **b**, CG MD simulations of the reentrant phase behavior. The top images illustrate the initial configurations of three simulation systems having, from left to right, 2, 8 or 16 RNA_40_ and 24 RP_3_ peptide molecules, realizing the 200 *µ*M peptide concentration used in experiment. The bottom row displays the configuration of each system after 2 *µ*s of CG MD simulation. The peptide molecules are shown in red, the RNA in blue. Blue squares illustrate the size of the simulation unit cell. **c**, Fraction of molecules forming the largest cluster in the simulations of the RP_3_–RNA_40_ mixtures differing by the number of RNA molecules. **d**, Average partition of molecules into the largest cluster as a function of the RNA-to-peptide weight ratio (red, the lines are guides to the eyes). The partitioning data were averaged over the last 3 *µ*s of each simulation. The blue line shows the behavior expected from the charge stoichiometry model Eq. (3) with the *x* axis defined at the top of the graph.

The observed reentrant phase behavior can be largely explained by the following charge stoichiometry considerations. Since one RNA_40_ carries a charge −40*e* (hereafter *e* denotes the charge of a proton) and one RP_3_ peptide carries a charge of +6*e*, 40/6 RP_3_ molecules are required to fully neutralize one RNA molecule, which corresponds to the RNA-to-peptide weight ratio of 1.2. Assuming the phase separation is driven by hydrophobic interactions between RP_3_ molecules and that it requires nearly complete neutralization of the electrostatic interactions, the largest cluster should be composed of RNA molecules each neutralized by 40/6 RP_3_ peptides. Then, the partitioning into the largest cluster has the following dependence on the number of RNA, *N*_RNA_, and peptide, *N*_pep_, molecules:

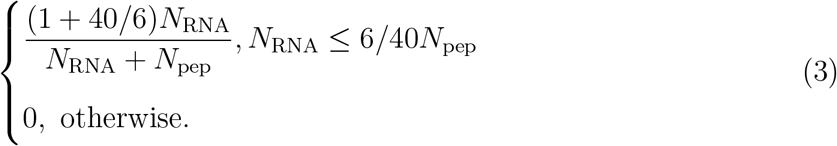

According to the charge stoichiometry model, no stable droplet can be formed when the charge of the RP_3_ peptides is not sufficient to compensate the charge of the RNA, i.e., when *N*_RNA_ *>* 6*/*40*N*_pep_. Despite its simplicity, the charge stoichiometry model reproduces the simulated reentrant behavior rather well, Fig. 2d, except near the peak of the dependence, where hydrophobic attraction is expected to dominate over pure electrostatic interactions.

### RNA alters the physical properties of a FUS condensate

The signature of a liquid-liquid phase transition is formation of stable droplet-shaped clusters containing proteins and nucleic acids surrounded by a dispersed phase where the proteins and nucleic acids are dissolved in solution. To obtain such stable two-phase coexistence and determine the effect of RNA on the properties of the condensate, we first simulated spontaneous phase separation in two simulation systems each containing 216 FUS proteins confined to a 136.5 × 136.5 × 136.5 nm^3^ unit cell. The simulations were performed at 292 K and one of the simulation systems additionally contained 84 copies of RNA_40_, Fig. 3a. That number of RNA molecules was chosen to approximately neutralize the electrical charge of the FUS proteins, with each protein carrying a charge of +15.5*e* according to the KH model. After 0.4 *µ*s of constant volume equilibration, a single droplet containing FUS or FUS and RNA molecules started to form, reaching equilibrium configuration by 1 *µ*s. That equilibrium configuration was then replicated to generate a 2 × 2 × 2 array of droplets, such that the nearest-neighbor droplets were separated by a 61 nm center-of-mass distance. Upon placing the eight-droplet arrays in a 273 × 273 × 273 nm^3^ unit cell, the two systems were simulated for another 400 ns, which was sufficient to observe complete fusion of the eight droplets into a single larger droplet. Movie 4 and 5 illustrate the fusion process in each system.

**Figure 3:**
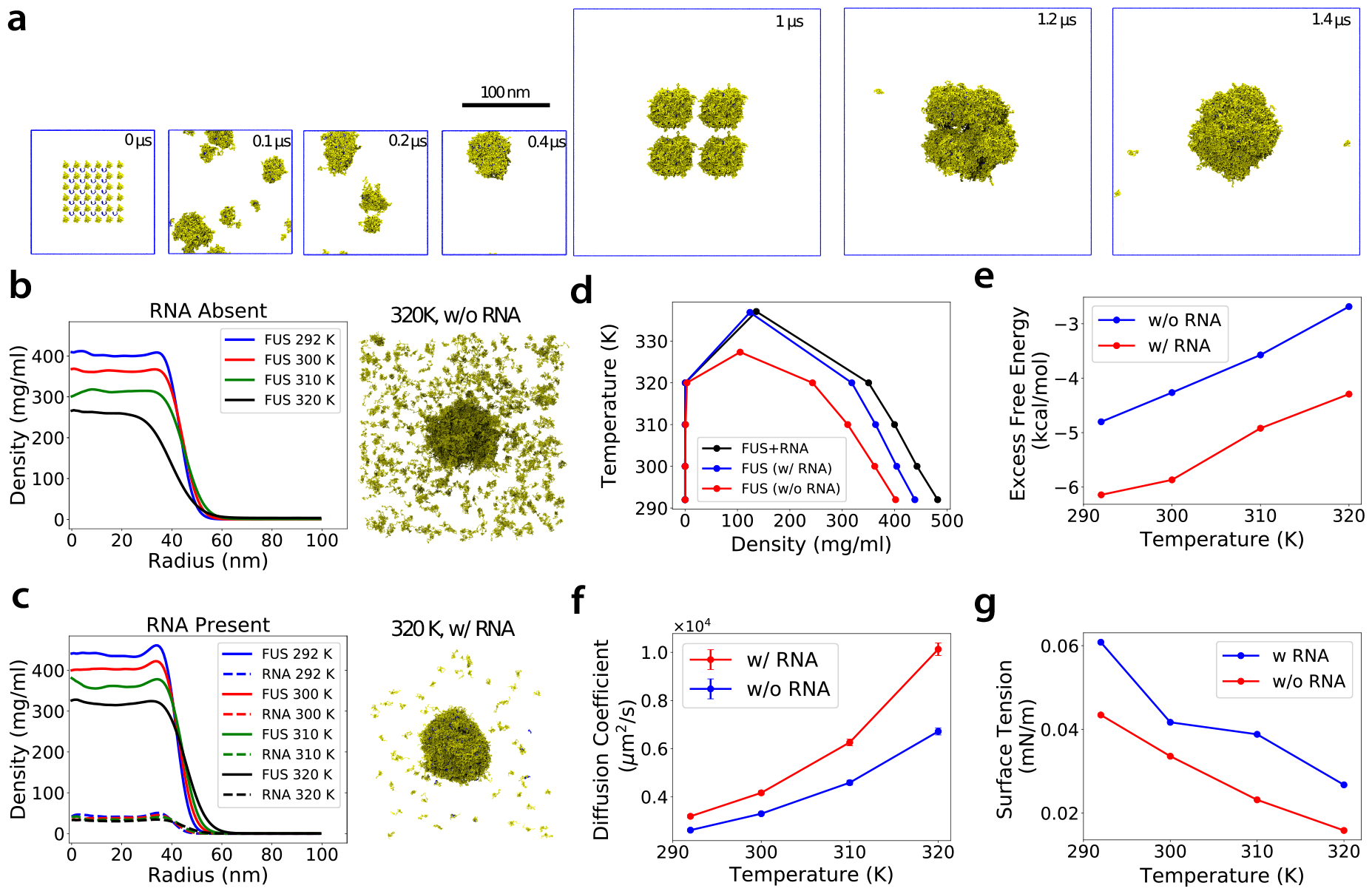
RNA alters temperature-dependent properties of a FUS condensate. **a**, Preparation of a condensate droplet containing 1728 FUS (yellow) and 672 RNA_40_ (blue) molecules. The droplet was constructed by first simulating spontaneous phase separation (at 292 K) of a smaller condensate system containing 216 FUS and 84 RNA_40_. The final configuration of the smaller droplet (after 1*µ*s) was replicated to form a 2×2×2 array and simulated until one large droplet was formed. This final configuration was used to initiate simulations at different temperatures. **b**, Radial density profile of a pure FUS condensate droplet at several temperatures. The image on the right shows a representative configuration at 320 K. **c**, Same as in panel b but for a FUS–RNA condensate droplet. **d**, Phase diagrams of pure FUS and FUS-RNA condensate systems. The blue and black curves both pertain to the system containing both FUS and RNA but differ by the density of species considered to construct the phase diagram (blue for RNA and FUS and black for FUS only). **e**, Excess free energy of a FUS molecule inside a condensate droplet at several temperatures, with and without RNA. **f**, Diffusion coefficients of a FUS protein within the condensed region estimated from the auto-correlation of the end-to-end distance. **g**, Surface tensions of a FUS droplet at several temperatures, with and without RNA.

After generating single-droplet systems containing either 1728 FUS molecules or 1728 FUS and 672 RNA_40_ molecules, we then gradually increased the temperature of each system from 292 K to another target temperature (300, 310 or 320 K) with the rate 0.1 K/ns. Upon reaching a target temperature, each MD system (including the 292 K one) was simulated at that temperature for additional 2 *µ*s and the first 500 ns were discarded from subsequent analysis.

Figure 3b plots radially averaged density of FUS in a pure FUS condensate system as a function of distance from the center of mass of the droplet. The FUS density is the highest at the lowest temperature (292 K) and decreases gradually as the system’s temperature increases. In the 292–310 K range, the width of the interface between the condensate droplet and the surrounding dispersed phase widens as the system’s temperature increases while the physical size of the droplet remains unchanged. At 320 K, though, the droplet not only has a substantially lower inner density, the droplet also shrinks in size, indicating substantial loss of FUS to the dispersed phase, as illustrated by the snapshot in Fig. 3b and Movie 6.

The presence of RNA moderately (up to 10%) increases the density of FUS within the RNA–FUS condensate and forces the condensate to retain its size within the whole range of temperatures studied, Fig. 3c. Interestingly, the FUS concentration is seen to reach a local maximum at the surface of the condensate, which is reminiscent of interface features previously seen in other complex coacervates with compositional asymmetries ^55,56^ At 320 K, substantially fewer FUS molecules are observed in the dispersed phased, Fig. 3c and Movie 7, in comparison to the pure FUS system, Fig. 3c, indicating stronger cohesion within the condensate. The density profile of RNA_40_ within the condensate generally mirrors that of FUS, with the ratio of FUS and RNA concentrations ranging between 9.9 and 10.4, depending on the temperature, which is close to the overall FUS-to-RNA weight ratio of 10.6.

The four sets of simulations allowed us to determine the effect of RNA on the phase diagram of FUS. For each temperature, we determined the average concentration of the condensed phase, *ρ*_*h*_, by averaging FUS density along the respective trajectories and within a 40-nm radius sphere centered at the center of the droplet. Similarly, we determined the average concentration of the disperse region, *ρ*_*l*_, by averaging over the volume extending beyond a 70-nm radius. Plotting the temperatures on the *y*-axis and the corresponding densities of the two phases on the *x*-axis, we obtained the phase diagrams of the two systems, Fig. 3d. The phase diagram of the pure FUS system (red) is consistent with that determined earlier^34^ using a slab-method. The two representations of the phase diagram for the FUS+RNA system (blue and black) differ by the choice of density used as the order parameter: the combined concentration of FUS and RNA (blue) and only of FUS (black). The critical temperatures, *T*_*c*_, for each phase diagram is found using the following relation^33,34^: *ρ*_*h*_ − *ρ*_*l*_ = *C*(*T*_*c*_ − *T*)^0.326^, where the critical exponent is that of the 3D Ising universality class^57^. The obtained critical temperatures were 327.3 K (red) for pure FUS condensate and 337.1 K (black) for the FUS+RNA mixture, with the latter critical temperature becoming only 0.2 K less if only FUS concentration (blue) is used as the order parameter. According to the rectilinear law, the critical density *ρ*_*c*_ is obtained by fitting the average density 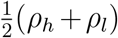 to *F* (*T* − *T*_*c*_) + *ρ*_*c*_, where *F* is a fitting constant. The critical density was found to be 123.8 mg/ml and 105.1 mg/ml for the FUS condensate with and without RNA, respectively. Thus, the presence of RNA increases the critical temperature of a FUS condensate by 9.8 K and extends the region of two-phase coexistence.

The snapshots of the two systems shown Fig. 3b and c along with Movies 6 and 7 clearly show that the condensate droplet containing RNA can more readily absorb molecules from the dispersed phase. To quantify the degree of “stickiness” of a condensate, we evaluated the excess free energy required to bring one FUS molecule into the droplet, i.e., −k_B_*T* ln(*ρ*_*h*_*/ρ*_*l*_), where *ρ*_*h*_ and *ρ*_*l*_ are the densities of FUS in the condensed and dispersed phases, respectively. Figure 3e plots the resulting excess free energy for both condensate systems as a function of temperature. Experimentally, the excess free energy for the LCD domain of FUS was found to range between 2 and 3 kcal/mol at 300 K^29,58–61^. The excess free energy decreased by ∼1.5 kcal/mol in the presence of RNA in the condensate, making it more favorable for the FUS molecules to become a part of a droplet. Considering together with the phase diagrams, Fig. 3d, we conclude that RNA increases propensity of FUS to droplet formation. This conclusion is in general agreement with experiments that found RNA to promote phase separation of FUS^15^ as well as with similar observations for other types of RNA-binding proteins^58,62,63^.

To investigate how RNA modulated the fluidity of a FUS droplet, we computed the diffusion coefficients of FUS inside the largest condensate droplet. When evaluating the auto-correlation function according to Eq. (10) (see Methods), we only considered those time intervals during which a FUS molecule continuously resided within the droplet. Figure 3f plots the resulting dependence of the diffusion coefficients on temperature with and without RNA in the droplet, along with the error bars estimated from the standard deviations of the relaxation times *τ*_1_ and *τ*_2_, see Methods for details. While in both systems, the diffusivity was found to increase with temperature, the presence of RNA made the condensate droplets more viscous. Such a decrease of fluidity upon RNA addition contrasts experiments that found RNA to increase fluidity of a LAF-1 condensate^13^, but are in qualitative agreement with another study that found RNA to decrease fluidity of a WHi3 condensate^64^. Our results are in line with a recent computational study that found RNA to increase the viscosity of a FUS–RNA condensate for RNA of comparable length ^41^.

Finally, we determined the surface tensions of the largest condensate droplet using the principal component analysis described in the Methods section. The surface tension was found to decrease with temperature regardless of the presence of RNA in the condensate, Fig. 3g. At each temperature, however, the presence of RNA increased surface tension making the droplet less deformable. Interestingly, side-by-side comparison of two trajectories illustrating the fusion of eight droplets into one larger droplet, Movie 4 and Movie 5, suggests that the increased by RNA surface tension of the condensate wins over its increased viscosity as the fusion results in a well-formed single droplet approximately four times faster in the system containing RNA.

### RNA length modulates cohesiveness and fluidity of FUS condensates

Experiments performed on different biological condensates showed that incorporation of RNA could have opposing effects on their physical properties, for example, increasing ^13^ or decreasing^64^ the condensate’s fluidity. One possible explanation of such puzzling behavior is that the effect of RNA depends on its relative length with respect to the spacing of the RNA-binding proteins within the condensate. To systematically investigate this possibility, we changed the length of the RNA molecules while keeping the total amount of RNA nucleotides in the condensate and, hence, the RNA-to-FUS weight ratio constant. Specifically, we constructed eight systems each containing 64 FUS proteins and either 200, 100, 67, 50, 33, 25, 12 or 6 RNA molecules each consisting of 5, 10, 15, 20, 30, 40, 80 or 160 nucleotides, which roughly makes the RNA+FUS system electrically neutral. As a control, one additional system was built to contain only 64 FUS proteins. Starting from a dispersed arrangements of the FUS and the RNA molecules, each system was simulated for 2 *µ*s at 292 K. The systems were observed to developed phase-separated morphology within the first 500 ns of the respective MD simulation, the last 1.5 *µ*s were used for subsequent analysis.

Having identified the largest cluster (droplet) in each simulation, we computed partitioning of FUS and RNA molecules into it by computing the number of FUS or RNA molecule within the cluster and dividing that number by the total number of either FUS or RNA molecules present in the system. While almost all FUS molecules were observed to partition into the largest cluster, partitioning of RNA molecules depended on their length, Fig. 4a. A large fraction of RNA molecules fewer than 30 nucleotides in length were found in the dispersed phase, not being a part of a long-lived droplet. The RNA partitioning into the condensate droplet increased abruptly when the RNA length exceeded 30 nucleotides. The average density of FUS within the condensate increases monotonically as the RNA molecules become longer, whereas the average size of the droplet decreased, Fig. 4b.

**Figure 4:**
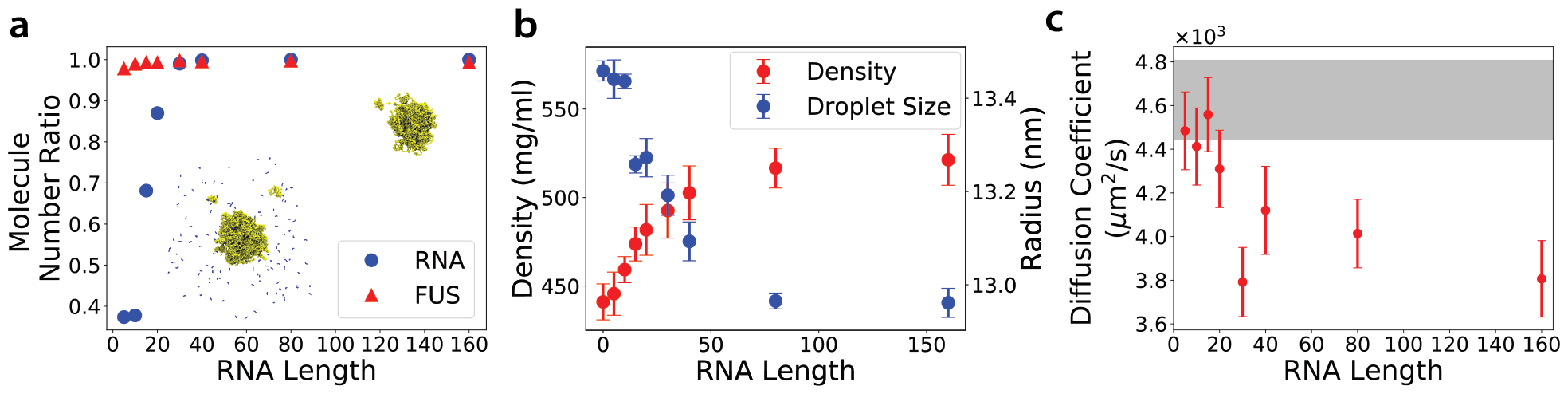
Effect of RNA length at constant RNA-to-FUS weight ratio on physical properties of a FUS condensate. **a**, Steady-state average fraction of FUS or RNA molecules (relative to the total number of either FUS or RNA) forming the largest cluster in CG MD simulations of eight systems each containing 64 FUS proteins and either 200, 100, 67, 50, 33, 25, 12 or 6 RNA molecules each consisting of 5, 10, 15, 20, 30, 40, 80 or 160 nucleotides, respectively. The RNA-to-FUS weight ratio is the same (0.24) in all eight simulations. The snapshots illustrates representative equilibrium configurations of the simulation systems containing RNA_10_ (bottom left) and RNA_160_ (top right). **b**, Average density of FUS within the condensate droplet (left axis) and the average radius of the droplet (right axis) as a function of the RNA length. The density was computed by averaging over the respective simulation trajectories and within a sphere of a 10 nm radius centered at the center of mass of the condensate. The error bars were estimated using the standard deviation of the densities inside the condensed region. The average size of the droplet was obtained by fitting the density profile to Eq. (11) (see Methods). The error bars for the size determination were estimated using the standard deviations of the fitted radii. **c**, Diffusion coefficient of FUS within the condensate as a function of RNA length. The region shaded in gray represents the range of diffusion constant values observed in the absence of RNA. All data were obtained at 292 K.

To characterize the effect of RNA on the condensate’s viscosity, we computed diffusion coefficients of FUS inside the droplet and compared it to the diffusion coefficient observed in the absence of RNA, Fig. 4c. We found that the presence of short RNA (*<* 20 nucleotides) barely alters the FUS viscosity, whereas longer RNA strands substantially reduce diffusion of FUS and thus increase viscosity of the condensate. Similar length dependence was reported from the study of LAF-1^17^, where the presence of short (*<* 30 nucleotides) RNA molecules was found to decrease the viscosity by a small amount while the presence of longer RNA strands increased the viscosity. Overall, we conclude that fluidity of a FUS condensate droplet decreases when more nucleotides are recruited into the condensate and that the effect becomes stronger for longer RNA molecules within the length range studied. These results suggests that long polynucleotides may play a role of a scaffold, binding at once to multiple RNA-binding domains, whereas short RNA molecules behave analogous to other small diffusive solutes permeating the droplet.

### RNA and temperature regulate porosity of a condensate droplet

The key property of membraneless organelles *in vivo* is their ability to recruit select classes of target molecules from the surrounding environment to locally enhance their concentrations and thereby enable a variety of function ^10,65^. Recruiting the molecules from the solution implies that such molecules are capable to enter and permeate through the condensate droplets. Dextran polymers of various length were used to experimentally probe diffusion of small molecules through LAF-1 droplets^17^, finding the diffusivity of the polymers to drop abruptly when their Stokes radius exceeds 3 nm. According to the free-volume theory ^66^, such a drop in diffusivity can be related to the drop of the free volume available inside the droplets for diffusing particles.

Visual inspection of the condensate morphology immediately reveals a sponge-like network of interconnected channels penetrating the condensate droplet. Filled with solution (represented implicitly in our simulations), such a network provides a path for smaller intruder molecules to enter and diffuse through the condensate. To quantitatively analyze such networks, we first charted the free volume inside the droplets by placing a sphere of a predetermined radius at the nodes of a 3D grid encompassing the condensate. The node is considered to be in free space if the sphere fits without clashes and occupied by the molecules of the condensate otherwise (see Methods and Ref. 67 for details). Figure 5a illustrates the free volume determined through such void analysis for three values of the probe radius: 0.4, 0.85 and 1.3 nm. As the probe radius increases, the free space inside the condensates becomes smaller in volume and less connected. The probe radius at which the connectivity of the free space disappears—the percolation threshold—should therefore mark a transition from a regime where the intruder molecules can diffuse unobstructed through the entire volume of the condensate to a regime where their motion through the condensates is governed by the rearrangements of the condenstate’s network, i.e., by the motion of the condensate molecules themselves.

**Figure 5:**
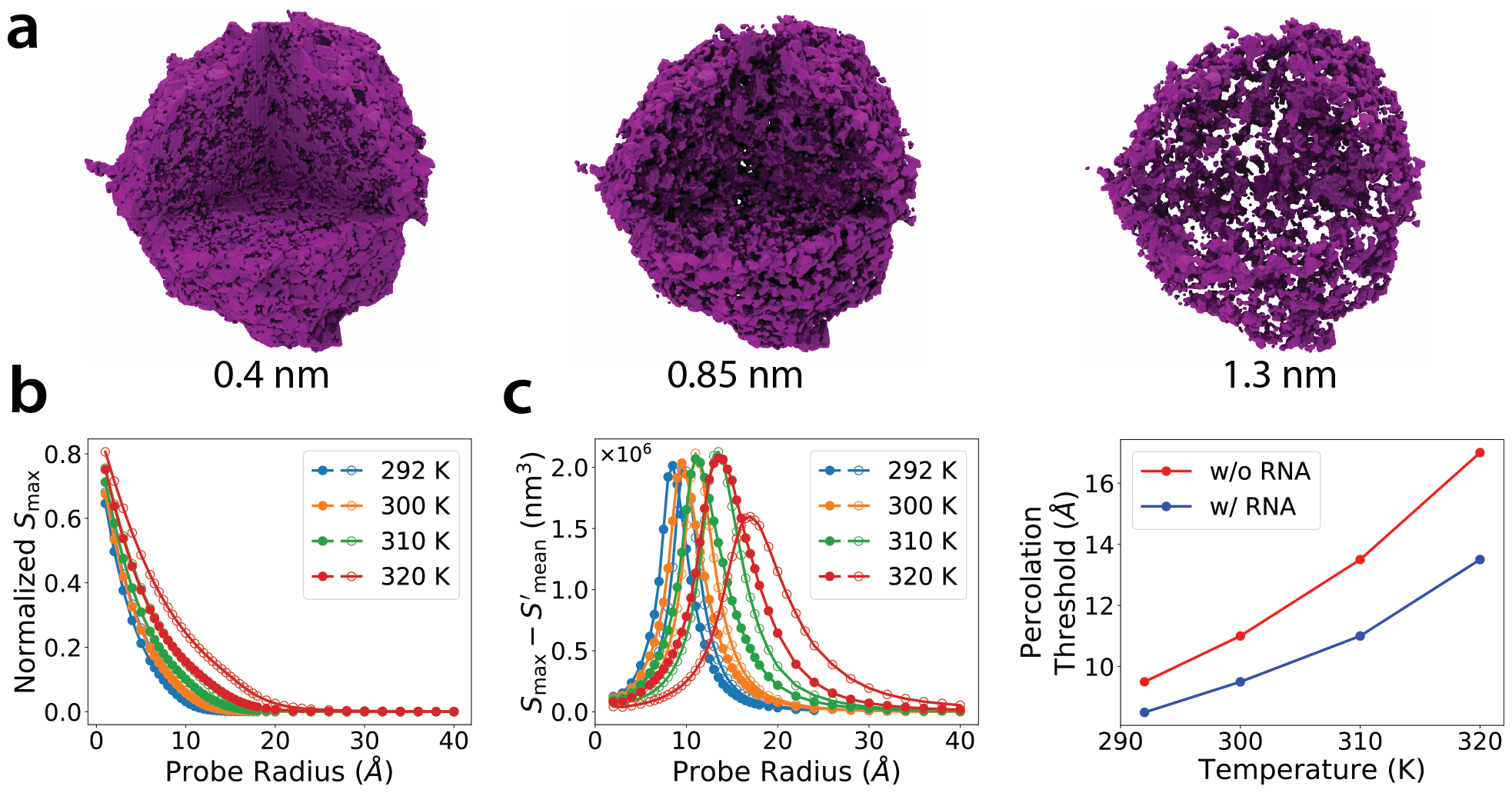
RNA and temperature regulate porosity of a FUS condensate. **a**, Void space, i.e., the space not occupied by the protein or RNA, within a condensate droplet obtained using a spherical probe of 0.4, 0.85 or 1.3 nm radius and visualized by filling the void volume with purple spheres. This particular instantaneous configuration of a FUS condensate droplet is taken from a CG MD simulation at 292 K performed in the absence of RNA. One eights of the condensate volume is cut away to better illustrate the internal network of channels within the condensate. **b**, The ratio of the largest cluster volume *S*_max_ to the total available volume as a function of probe radius at several temperatures. This ratio serves as the order parameters for the percolation transition. The filled and empty symbols denote simulations carried out with and without RNA, respectively. **c**, The difference between the largest cluster volume and the mean cluster volume, 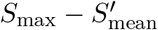, as a function of the probe radius at several temperatures. The difference attains a maximum at a percolation transition. **d**, The percolation transition radius as a function of temperature. The presence of RNA lowers the threshold by compacting internal voids of the droplet.

We find the percolation threshold by treating percolation as a geometrical transition characterized by the divergence of the largest connected cluster, which mathematically represents a merger of all small clusters into one large cluster upon the transition. Inside a droplet, clusters formed by free volume at different probe radii display a similar transition: under certain probe size, *p*_*c*_, the available free volume expands to entire droplet and forms the interpenetrating channels. Figure 5b plots the ratio of the largest free space cluster volume, *S*_max_, to the entire empty space volume in the droplet as a function of the probe radius for the condensate droplet containing 1782 FUS molecules and either zero or 673 RNA_40_ molecules at several temperatures (the same systems are featured in Fig. 3). As expected, the largest cluster volume decreases as the probe radius increases, reaching zero at approximately 2 nm. To quantitatively identify the percolation threshold, we plot in Fig. 5c the quantity 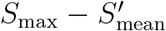, where 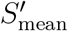 is the mean cluster volume obtained by averaging over all clusters. The maximum of that quantity indicates the onset of the percolation transitions, see Methods for details. Collecting all peak locations from Fig. 5c, we obtain the relation between the percolation threshold, temperature, and the presence of RNA in the condensate, Fig. 5d.

Through our void analysis, we thus find the probe radius at which the empty volume clusters no longer form a continuous path through the condensate droplet to increase with temperature, raising by more than twofold from 292 to 320 K. The presence of RNA, however, is found to considerably reduce the size of the channels penetrating the condensate. Considering that the volume occupied by RNA in the condensate is *<* 5% of the total volume, such a decrease of the percolation thresholds indicates that FUS molecules themselves are packed more tightly within the droplet in the presence of RNA. At 320 K and in the absence of RNA, the percolation threshold radius is found to be 1.7 nm, which is large enough to allow unstructured RNA molecules to enter and diffuse through the condensate. The compaction of the internal channels observed in the presence of RNA thus may serve as a physical mechanism limiting the entrance of additional RNA molecules into the condensate and thus the internal RNA concentration.

## Discussion

In this study, we demonstrated a systematic top-down approach for developing a custom CG model of RNA compatible with the KH CG model^32^ representation of a specific protein, FUS. Our RNA model is shown to have the persistent and contour lengths in close agreement with experiment and to reproduce the reentrant phase behavior of a peptide–RNA mixture. Importantly, our custom RNA model is parametrized to reproduce experimentally measured affinity of the RNA–FUS interactions.

Using our RNA model, we performed extensive computational investigation of FUS–RNA condensate droplets each containing approximately 2,500 individual molecules and spanning ∼ 0.1 *µ*m in diameter. We found the presence of 40-nucleotide RNA molecules to enhance propensity of FUS to form condensate droplets by decreasing its excess free energy. At the same time, the RNA is found to make droplets less fluid by increasing the condensate’s viscosity and surface tension. These results are in general agreement with the previous experimental studies of FUS ^15^ and other RNA-binding proteins^13,58,62–64^ carried out at low RNA concentrations.

To gain insight into the internal structures of the condensate, we devised an algorithm to identify the clusters formed by the free volume inside the droplets and determined the upper limit for the size of the probe molecules that could diffuse through such internal channels unobstructed. We observed the mesh size to shrink dramatically in the presence of RNA, suggesting that RNA compacts the droplets by forming inter-molecular networks. To test this hypothesis, we varied the length of the RNA strands while maintaining the same number of RNA nucleotides within the condensate. We observed a monotonic increase of the condensate concentration with the RNA length up to approximately 80 nucleotides. This behavior suggests that longer RNA strands indeed make the droplets adopt a denser structure. This conjecture is further supported by our measurements of FUS diffusion within the droplets: short RNA molecules barely affect FUS diffusivity in contrast to longer strands that reduce it. Combining all observations, we conclude that RNA strands exceeding 20 nucleotides in length serve as a scaffold of a condensate droplet, forming and stabilizing the inner network structures through binding to multiple inter-molecular RBD segments of FUS.

## Methods

### MD simulations of FUS-RNA condensates

All MD simulations were performed using the LAMMPS software package^68^, periodic boundary conditions and a 10 fs simulation time step. The temperature was kept constant using the Nöse-Hoover thermostat ^69,70^ and the default parameters of LAMMPS. A 3.5 nm cutoff was used for the calculations of all nonbonded forces, including the screened Coulomb electrostatics.

Our CG model of full-length FUS utilizes the parameterization of the KH model^32^, as described previously^34^. Briefly, each amino acid of the protein is represented by one CG bead of the corresponding mass. The beads are linked into a string according to the amino acid sequence of FUS^48^ via harmonic bond potentials of 3.8 kcal/(mol Å^2^) spring constant and 3.8 Å equilibrium length. The beads forming the two structured domains of FUS, i.e., the RNA recognition motive (residues 278 to 385) and the zinc finger domain (residues 419 to 454), are restrained to maintain their equivalent crystallographic distances ^48^ using harmonic potentials (3.8 kcal/(mol Å^2^) spring constant) that apply to any pair of residues having an equilibrium bead-to-bead distance between 8 and 12 Å. The electrostatic interaction between any non-neighboring charged beads is described within the Debye-Hückel approximation:

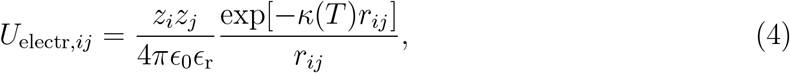

where *z*_*i*_ and *z*_*j*_ are the charges of residues *i* and *j* (*e* for arginines and lysines; −*e* for aspartic and glutamic acids and 0.5 *e* for histidines, where *e* is the charge of a proton), and *r*_*ij*_ is the distance between the residues. Relative permittivity *ϵ* _r_= 80 (the value of water), and the temperature-dependent screening length, 1/ *κ*(*T*), varies between 7.8 and 10.2 Å in the 292 to 320 K temperature range. In addition, each non-neighboring residue pair, (*i, j*), interacts via the following nonbonded potential

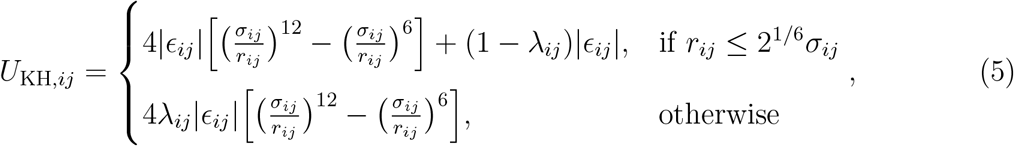

where *σ*_*ij*_ = (*σ*_*i*_ + *σ*_*j*_)*/*2 and the parameters *σ*_*i*_, *σ*_*j*_ and *ϵ*_*ij*_ were taken from the KH model^32^. The hydrophobicity parameter *λ*_*ij*_ is set 1 when *ϵ*_*ij*_ is negative and −1 otherwise.

In our custom CG model of RNA, each nucleotide is represented by one CG bead of 324.2 amu mass and electrical charge of −*e*. Neighboring beads are connected via harmonic bond potential

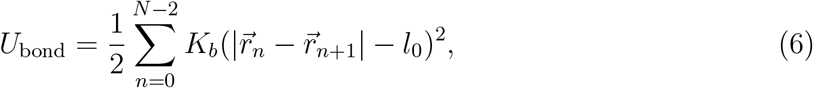

where *K*_*b*_ = 10 kcal/(mol Å^2^) and the equilibrium bond length *l*_0_ = 5.036 Å. A valence angle between each three neighboring beads, *P*_*n−*1_, *P*_*n*_, *P*_*n*+1_, is subject to an angle potential

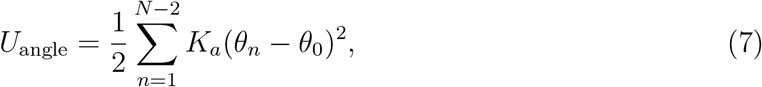

where *K*_*a*_ = 2 kcal/(mol degree^2^), *θ*_*n*_ is the instantaneous value of the valence angle between the three beads, and *θ*_0_ = 141.8 degrees is the equilibrium valence angle value. The specific values of the bond length and the valence angle were determined by fitting small angle x-ray scattering (SAXS) data, as described in Results.

In addition to the bond and angle potentials, CG beads representing two nucleotides (excluding neighbors and next nearest neighbors) interact via the Weeks–Chandle–Andersen (WCA) potential, a variant of the Lennard-Jones potential,

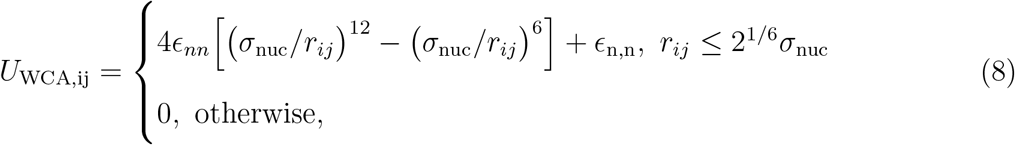

where the van der Waals radius, *σ*_nuc_, of each RNA nucleotide is 5.58 Å and the interaction strength *ϵ*_n,n_ = 0.58 kcal/mol. The same two nucleotides also interact electrostatically according to Eq. 4.

The nonbonded potentials describing the interactions between amino acids of FUS and the RNA nucleotides have both electrostatic and Lennard-Jones contributions. The electrostatic interactions are calculated using Eq. 4 and the nominal charges of the amino acids and the nucleotides. The Lennard-Jones interactions are calculated using Eq. (5) and two custom energy scale parameters: *ϵ*_n,d_ = 0.001 kcal/mol that describes interactions between an RNA nucleotide and any unstructured amino acid of FUS and *ϵ*_n,s_ =−0.131 kcal/mol that describes interactions between an RNA nucleotide and any amino acid of any structured domain of FUS. The specific values of the parameters are derived and validated as described in Results.

### Potential of mean force (PMF) calculations

The FUS–FUS and FUS–RNA PMFs were computed using the adaptive bias force (ABF) method^51^ implemented in LAMMPS as a COLVARS module^51,71,72^. To ensure adequate sampling of the PMF, the whole range of the reaction coordinate was split into a number of overlapping sections, the number of sections varied between six and twelve, depending on the length of the RNA strand. The width of each window was 5 nm and the overlap between the neighboring windows was 3 nm. The simulations in each window lasted 4 *µ*s and were carried out at 292 K. The reaction coordinate was constrained to remain within the desired range in each window by means of two half-harmonic potentials with 20 kcal/(mol Å^2^) spring constant. The PMF was obtained by merging all the samplings at each window using the protocol implemented in LAMMPS.

### Diffusion coefficient

Following the method described by Hummer^73^, we estimated the diffusion coefficient as

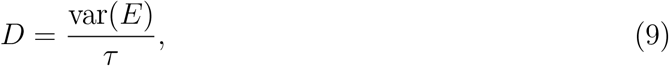

where var(*E*) is the variance of the end-to-end distance *E* and the characteristic time *τ* is estimated from the auto-correlation function *δE*(*t*) ≡ *E*(*t*) − ⟨*E*⟩,

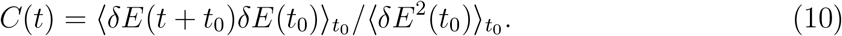

Above, 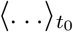 indicates an average over all possible initial times, *t*_0_, along an MD trajectory. The auto-correlation function *C*(*t*) was fitted by a biexponential expression *A* exp(−*t/τ*_1_) + *B* exp(−*t/τ*_2_), following which *τ* were determined as a weighted average (*Aτ*_1_ +*Bτ*_2_)*/*(*A*+*B*), see Ref. 29 for details.

### Cluster analysis

Two molecules were assigned to the same cluster if any intra-molecular pair of their residues was separated by less than the sum of the residues’ van der Waals radii. Our search for neighboring particles utilized conventional cell list data structure^74^. The volume of the simulation cell was split into cubic cell, 2 nm on each side, which was large enough to accommodate all possible pairs of residues that form a cluster contact according to our definition. After identifying all contact pairs for every particle, any two molecules having at least one contact pair were marked as belonging to the same cluster. A disjoint-set data structure^75^ was then used to represent clusters formed by the condensate molecules.

### Radial density profile

When estimating the radial density profile, the center of mass of the largest cluster was shifted to the origin and the radial density with respect to the origin was computed. The average radius of the droplet denoted as *R* was extracted by fitting the radial density profile to the following relation,

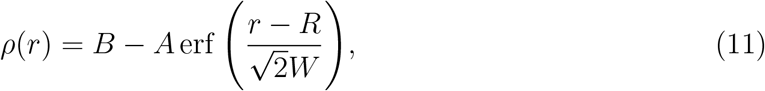

where *A* and *B* are two fitting constants and *W* is the average width of the interface.

### Surface tension

Surface tension of a condensate droplet was determined following a method proposed by Henderson and Lekner^76^, which was recently used for analysis of biological condensates^29^. Thermal fluctuations of a droplet produce changes of its surface area, *δA*, which are related to the surface tension, *γ*, as *U* = *γδA*, where *U* is the potential energy stored at the interface. Following the derivation presented in Ref. 29,

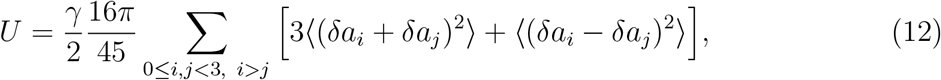

where *δa*_*i*_ ≡ *a*_*i*_ − *R* is the fluctuation along a principal axis *i* with respect to the average radius *R* obtained using the fitting approach described above. According to the equipartition theorem, each of the two modes ⟨ (*δa*_*i*_ + *δa*_*j*_)^2^⟩ and ⟨ (*δa*_*i*_ − *δa*_*j*_)^2^⟩contributes 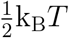 to the potential energy. Hence, the surface tension estimates for the two modes are

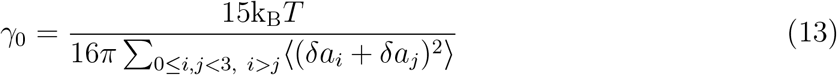

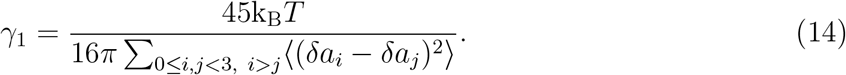

We determined *γ*_0_ and *γ*_1_ by computing, for each frame of a CG MD trajectory, a 3×3 weighted covariance matrix

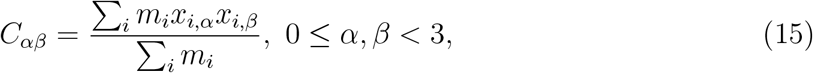

where *x*_*i, α*_ is the *α*-axis coordinate of a CG particle *i* (nucleotide or amino acid) and the sum runs over all molecules forming a droplet. The matrix was diagonalized to obtain *λ*_0_, *λ*_1_ and *λ*_2_ as its eigenvalues, from which 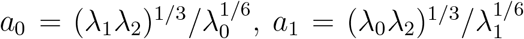 and 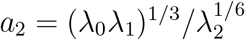. Knowing the trajectory-average radius of the droplet *R*, Eq. 11, instantaneous values of *δa*_*i*_ were obtained. The procedure was repeated to obtain trajectory-average values of *δa*_*i*_ and, subsequently, *γ*_0_ and *γ*_1_, which were within 1% of one another.

### Percolation transition

Upon identifying the largest cluster (droplet) of the condensate in an MD trajectory, each frame of the trajectory was translated to place the center of mass of the largest cluster at the origin. The system then was discretized in 2 Å bins along each dimensions. Each cubic cell voxel was represented by the coordinates of its center; only voxels inside the cluster were considered in the subsequent void analysis.

To identify the interior voxels, we use the *α*-shape procedure^77^ to define the shape of the cluster, *i*.*e*., to construct a polyhedron enclosing all protein and nucleic acid residues of the cluster. In our implementation, we took advantage of the Computational Geometry Algorithms Library (CGAL)^78^ to construct the *α*-shape and to choose the *α* value that produced only one connected component enclosing the cluster.

For each voxel located within the condensate cluster, we determined the largest sphere of radius *p* and the origin at the cluster that fits within the condensate without overlapping with any particles of the condensate. The radius of each particle was set to half of the particle’s vdW radius, *σ*. For a probe of a given size, *p*_probe_, we classify voxels as available if *p > p*_probe_ and occupied otherwise. Any two neighboring available voxels were assigned to the same cluster.

The order parameter for a percolation transition is the size of the largest cluster, *S*_max_, which diverges for an infinite systems when *p*_probe_ approaches a percolation threshold, *p*_c_. The mean cluster size *S*_mean_ is given by

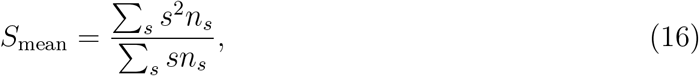

where *n*_*s*_ is the number of clusters having *s* voxels. *S*_mean_ is the expectation size of a cluster to which a randomly chosen available voxel belong to. Note that, Eq. (16) excludes the cluster containing infinite number of voxels and diverges for an infinite system when *p*_probe_ approaches a percolation threshold, *p*_c_. For a finite system, however, direct evaluation of Eq. (16) is not straighforward as the value of *p*_c_ is not known *a priori*^79^. Followin the algorithm introduced in Ref 79, we instead evaluated the quantity, 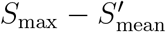 to detect the percolation threshold, where 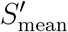 is the mean cluster size obtained by averaging over all clusters. The peak of the 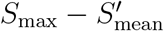 function indicates the percolation threshold.

## Acknowledgements

This work was supported by the National Institutes of Health through grant R01-GM137015. The supercomputer time was provided by the ACCESS allocation MCA05S028, and Leadership Resource Allocation MCB20012 on Frontera at the Texas Advanced Computing Center. Frontera is made possible by National Science Foundation award OAC-1818253.

## Author contributions

A.A. conceived the study. H.Y.-C. developed the CG model of RNA; performed all simulation and analysis. Both authors contributed to writing and editing of the manuscript.

